# Preserving Information and Integrity in Cryo-EM: A Fourier-Space Deposition Approach

**DOI:** 10.1101/2025.11.26.690836

**Authors:** Zbyszek Otwinowski, Yirui Guo, Raquel Bromberg, Dominika Borek

**Affiliations:** Department of Biophysics, The University of Texas Southwestern Medical Center, 5323 Harry Hines Blvd., Dallas, TX, 75390, USA; Department of Biochemistry, The University of Texas Southwestern Medical Center, 5323 Harry Hines Blvd, Dallas, TX, 75390, USA; Ligo Analytics, 2207 Chunk Ct., Dallas, TX, 75206, USA

**Keywords:** cryo-EM deposition, Wiener filter, half-map Fourier Shell Correlation, molecular masks, effective resolution

## Abstract

Current guidelines for depositing cryogenic electron microscopy single particle reconstruction (cryo-EM SPR) data require submission of unfiltered, unmasked, and unsharpened raw half-maps. The Fourier Shell Correlation (FSC) between the half-maps is then used as a proxy for the signal-to-noise ratio (SNR) to estimate the reconstruction’s resolution. This policy was introduced to enable independent validation of reported resolutions. Although developed to safeguard data integrity and minimize bias, these guidelines do not account for specific features of modern cryo-EM processing software, in particular weighting schemes that are not retained in half-map depositions and yet in general are necessary to recapitulate resolution estimates. As a results, resolution estimates and other validation statistics based on half-maps FSC may be under- or overestimated. Here, we describe the limitations of the current deposition guidelines and propose an alternative: depositing cryo-EM results in Fourier (reciprocal) space together with the mandatory deposit of molecular masks or their descriptors. This approach addresses the current limitations, preserves critical information from the reconstruction process, and better supports downstream analyses.

**Highlights:**

- Reciprocal-space deposition to preserve signal and uncertainty estimates.
- Mandatory molecular masks deposition for accurate and unbiased FSC validation.
- Limitations of half-map-based Fourier Shell Coefficients.
- The difference maps can be calculated.

*Graphical abstract:* 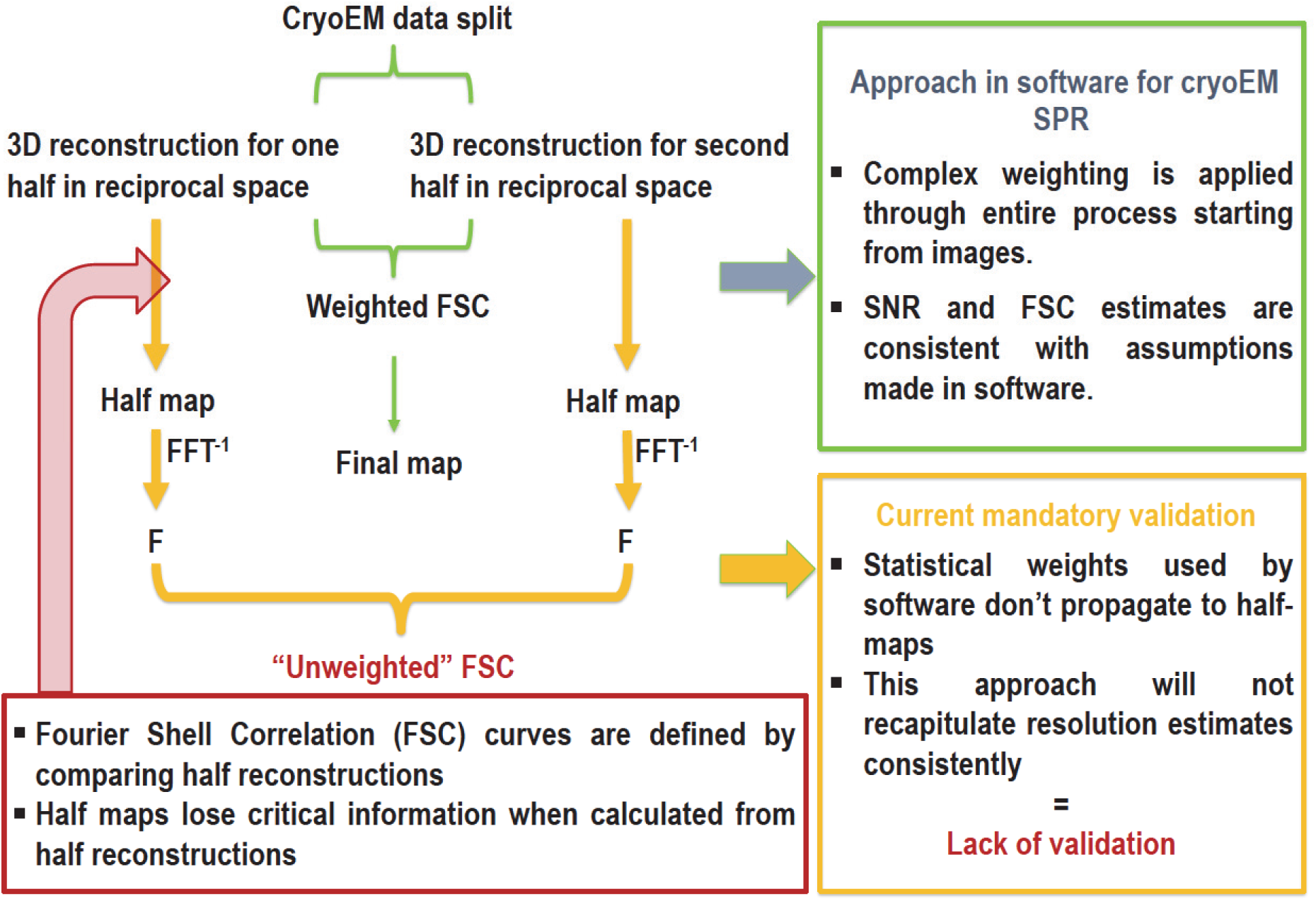

## Introduction

The Fourier Shell Correlation (FSC) between half maps is a core statistical measure in single particle reconstruction (SPR) and is used to calculate signal-to-noise ratio (SNR) statistics^1^.

The resulting SNR estimates are affected by how masking is applied and propagated in these calculations, with significant variability both within and between programs. Masks may be explicit when derived directly from 3D reconstruction, or implicit^2^, when only the mask volume is estimated from particle’s molecular weight. Thus, interpretation of FSC between half maps cannot be considered without knowing the specifics of masking process^3^.

There are several misconceptions regarding masking impact on half-maps FSC calculations in iterative 3D reconstruction. Although, the Protein Data Bank (PDB)^4^ requires that deposited model is accompanied with deposition of reconstructed maps to the Electron Microscopy Data Bank (EMDB)^5, 6^ and include “mandatory half-maps [that] must be unfiltered, unmasked, unsharpened, and positioned in the same coordinate space and orientation as the primary map such that they superimpose” (Figure 1), deposition of masks used in the reconstruction is encouraged but not mandatory^7^. The current standard was introduced in 2022 based on a recommendation from the 2020 wwPDB single-particle cryo-EM data-management workshop^7^. The requirement for depositing minimally-processed half maps reflects two underlying motivations: a concern that masking may introduce bias^8^ and the desire of repositories such as the PDB and EMDB to compute FSCs internally in order to validate the reported resolutions of deposited maps independently. However, the requirement to deposit unfiltered, unmasked, unsharpened half-maps creates several problems.

**Figure 1.**
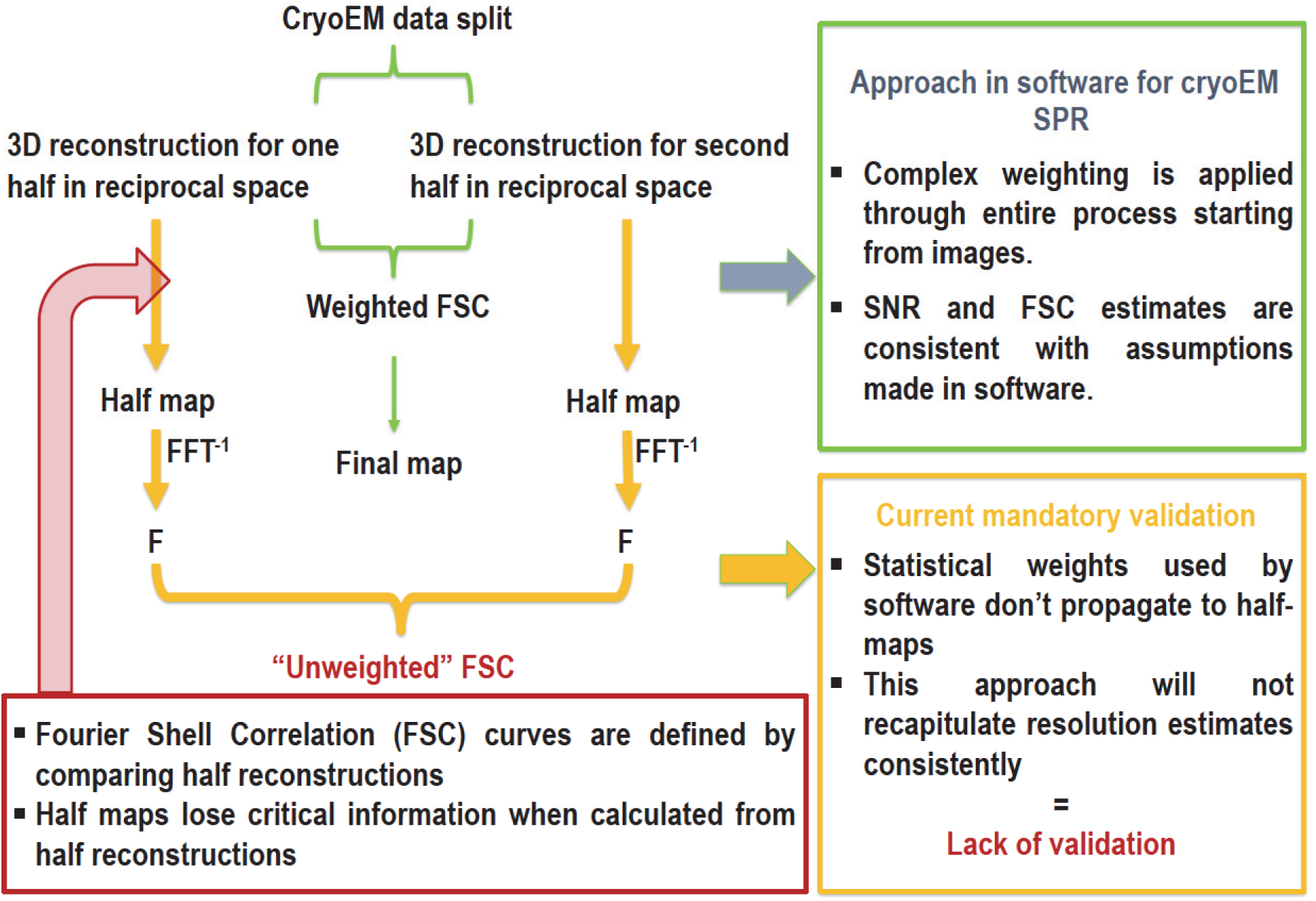
The half maps use under the current deposition rule. “Unfiltered” half maps do not contain information about weighting used during reconstruction. Because weighting is applied throughout 3D SPR to optimize resolution, the absence of these weights means that the resolution estimates reported by depositors cannot be reproduced.

FSC calculations starting from deposited unfiltered half maps may or may not reflect the true resolution of the reconstruction. FSC calculation from unfiltered half maps may be useful when a dataset shows no preferred orientation and when mask is used during the FSC calculation. However, in the case of strong preferred orientation, FSC calculation has little to no meaning (Figure 2), because information about local coverage in Fourier (reciprocal) space is missing. Local coverage is defined by how many planes of particle FFTs contribute to a particular point in the Fourier space, e.g., sum of CTF^2^ during back projection in cisTEM^2,9^ or Posterior Precision Directional Distribution in CryoSPARC^10^. During an actual reconstruction calculation, the local reciprocal space coverage information is used to calculate the Wiener filter. The Wiener filter can then be used to generate filtered half maps, which the PDB relies on to compute the FSC. If only unfiltered maps are deposited, the PDB cannot reproduce this reconstruction step unless the reconstruction weights given by the sum of CTF^2^ at each reciprocal-space point are also deposited. The lack of CTF^2^, particularly in respect to angular dependence, is a serious shortcoming of the deposition process, because this information is not only needed for validation but also contributes to reciprocal space uncertainty estimates, which can be used in the atomic model refinement.

**Figure 2.**
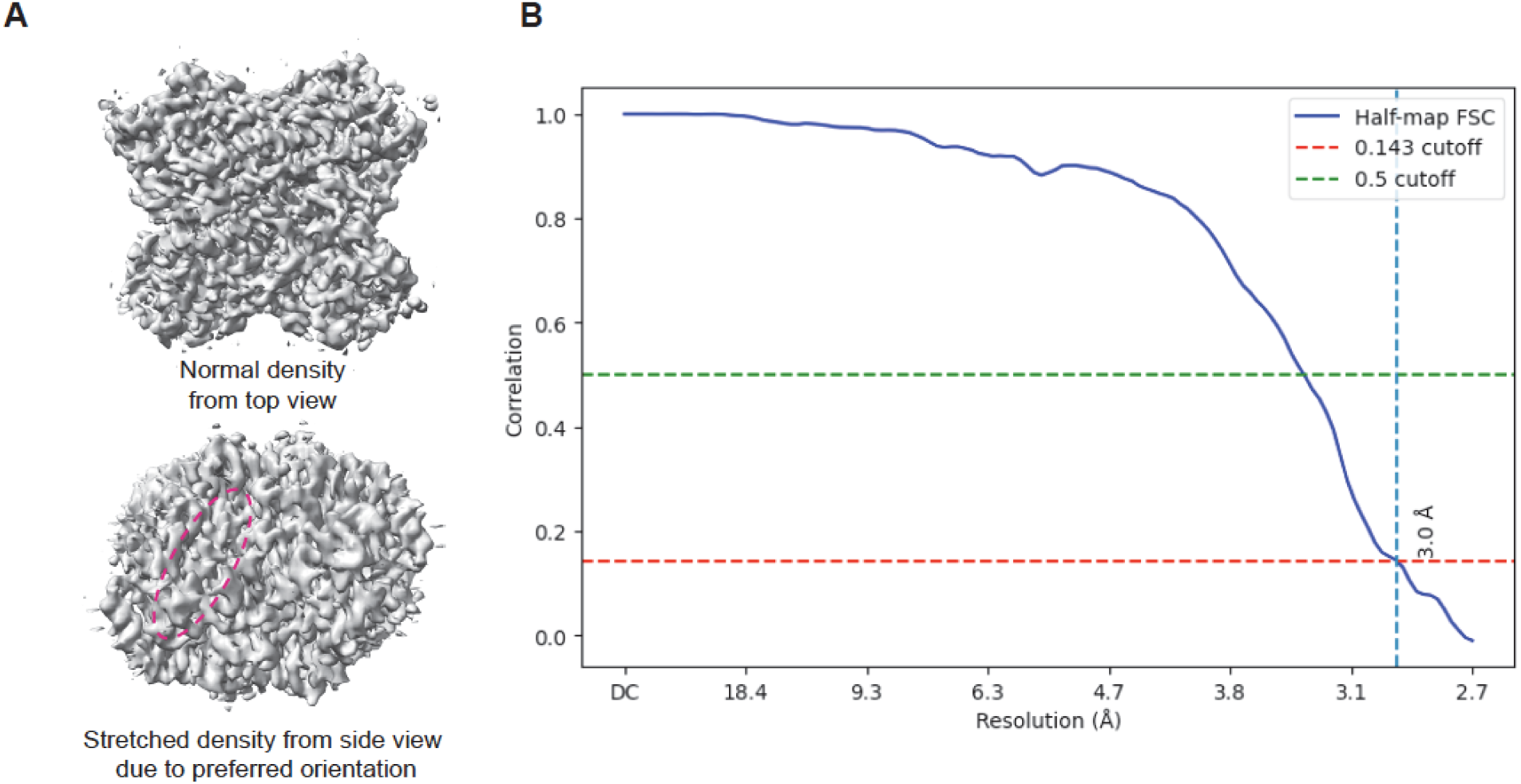
Glucose Isomerase (GI) reconstruction in the presence of strong preferred orientation showing a high half-map FSC. (A) Two views of the GI map reconstructed from particles with strong preferred orientation. The signature of preferred orientation is visible as stretched density, highlighted by the red dashed circle. (B) The FSC indicates resolution of 3.0 Å resolution. This is an example of overestimating map resolution. The 3.0 Å is real but does not extend uniformly across the full reconstruction sphere.

Without applying masks, the EMDB-recalculated FSC on half maps severely underestimates the resolution (Figure 3). The figure shows that the FSC values based on deposited half maps do not agree with the estimate provided by the depositors, but the agreement between model and data validates the depositor’s estimate.

**Figure 3.**
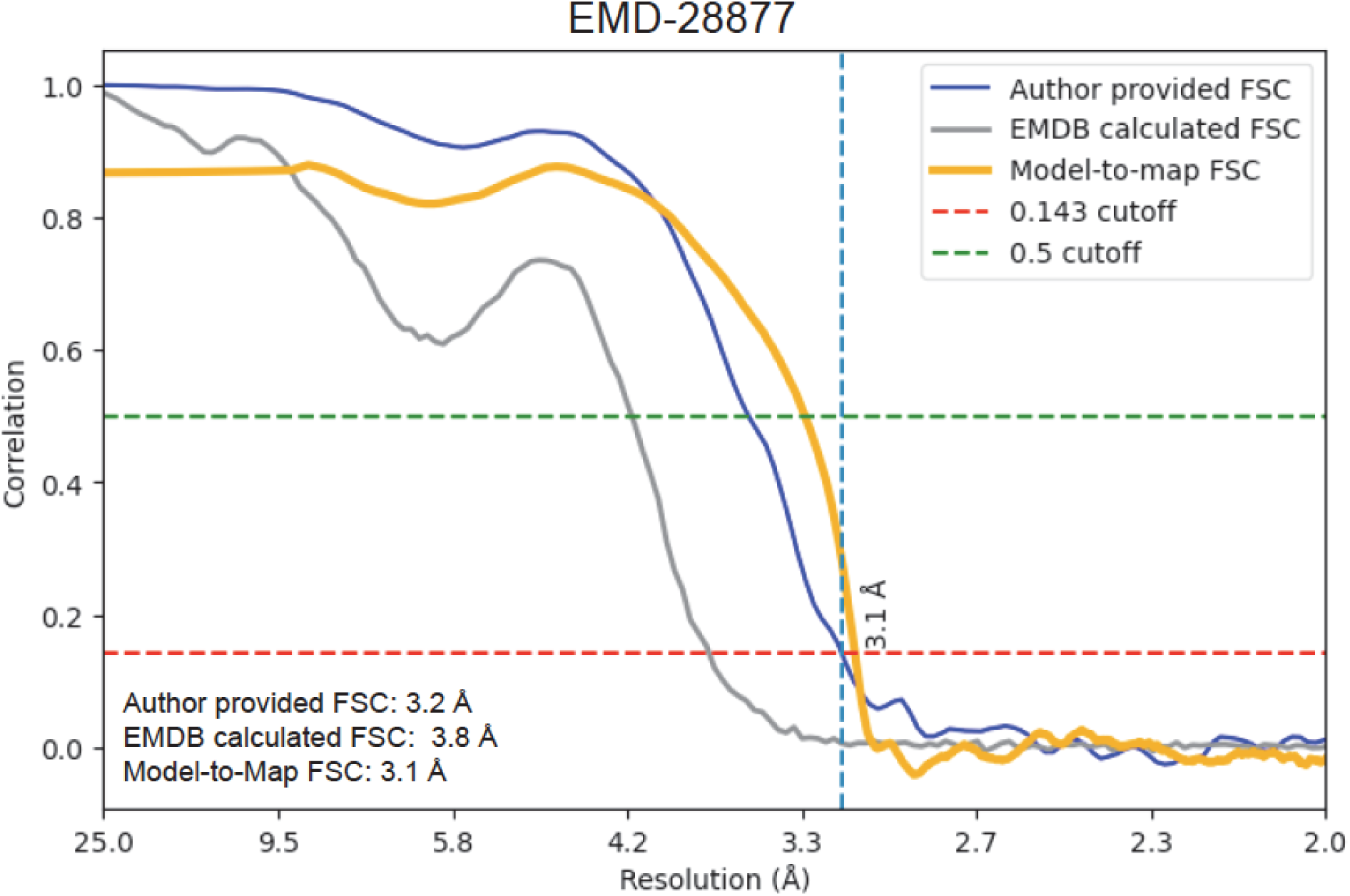
Example of the current PDB validation for EMD-28877^24^. The FSC calculated by the PDB from the deposited half maps (gray) does not the resolution estimate provided by the depositors (blue), whereas the model-to-map agreement (calculated with Phenix^3^) supports the depositor’s estimate (orange). See also examples in Wang, *et al*^25^.

Finally, the PDB requires that half maps must not be sharpened. One might expect this requirement to improve comparability across software packages (Supplementary Materials). However, differences in reconstruction algorithms can lead to significant variation in overall B-factor, even when the same particles are processed. For instance, a refinement in RELION may yield a different unsharpened B-factor compared to cryoSPARC, even if both reconstructions have properly converged. After post-processing (e.g., sharpening), the outputs from both programs often become similar. This indicates that the overall B-factor is not solely a property of the data, but also of the reconstruction procedure itself (e.g., masking strategies and whether angular marginalization was used). As a result, the restriction on sharpening does not effectively solve the problem of comparability between deposited raw maps. In fact, applying appropriate sharpening can help normalize differences introduced by various reconstruction workflows and make results more comparable across different software.

In short, the current deposition requirements pertaining to the submission of unfiltered, unmasked, and unsharpened half maps, have led to a disconnect between the EMDB-recalculated FSC and the actual signal-to-noise ratio (SNR) present in the final, processed map. The question therefore becomes how to preserve information from the reconstruction during the deposition and when to use FSC as a source of SNR estimation. This is especially important in reconstructions affected by preferred orientation. In such cases, FSC calculated from unweighted half maps will severely underestimate the quality of the reconstruction, while FSC from Wiener-filtered/weighted maps will overestimate it. FSC by its nature has no directional information and each shell is ascertained by a single number which is insufficient to represent maps that have directional quality, for example missing cone of data.

In addition, the meaning of “unfiltered, unmasked, unsharpened” becomes ambiguous as novel methods emerge, e.g., local refinement, where several filtering or masking technique serve as core aspects of reconstruction. Wiener filtering performs resolution-dependent calculations in reciprocal space and masking filters out noise in real space (Figure 4). The nature of the reconstruction process also implies that at least a circular mask should be used to provide anti-aliasing filter for particles from different angles inside the reconstruction box (Figure 4). It is likely the programs are already producing their intermediate calculations with filtered half maps.

**Figure 4.**
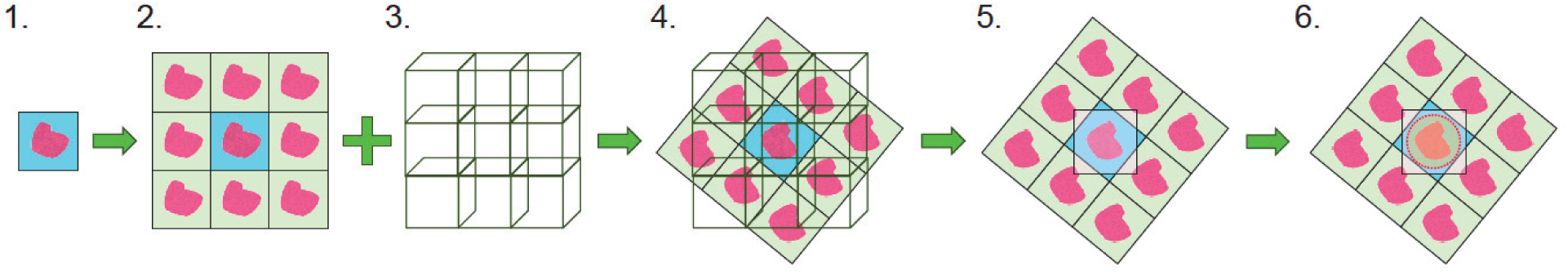
Reconstruction has to be masked. Steps 1-6 shows steps in typical reconstruction. 1. Particle image; 2. Set periodic boundary condition (assumption of FFT); 3. Reconstruction box with 3D periodic boundary condition; 4. Particle aligned to reconstruction box; 5. Schematic representation of intersection between two periodic boundary conditions. Corners of reconstruction box cannot be reconstructed due to aliasing; 6. Antialiasing filter has to be used. These reconstruction steps indicate that every reconstruction is masked. The 3D reconstruction is performed in the final volume. The information in this volume is propagated from the particle boxes that implicitly define a mask, as they are cutouts from the original micrograph. Many methods apply either explicit radial soft masks or implicit ones at the 3D accumulation stage (addition of all signals together), e.g. Kaiser-Bessel filtering or Gaussian filtering. Every approach to 3D reconstruction creates a spatial distribution of noise and understanding features of that noise distribution is necessary for estimating and validating resolution claims.

To enable clearer data-quality assessment and preserve directional information of the cryo-EM SPR reconstruction, we propose depositing reconstructions in Fourier (reciprocal) space, analogous to X-ray crystallography deposits. This format would preserve the original signal, its uncertainty, and the Wiener weights at each reciprocal-space point. The deposition of the molecular mask used for FSC calculations should become mandatory, with the masks deposited in the form used in the reconstruction process, i.e. either explicit (map file) or implicit (mask volume value). This approach allows transparent validation with unbiased resolution assessment, and appropriate treatment of preferred orientation and non-uniform sampling in reciprocal space. It also supports new methods for model refinement and would make deposits more compact in size. An additional motivation is preventing unintended or deliberate map’s alterations. Reciprocal-space deposition is more tamper-resistant than current deposits, because real-space edits cannot be matched by consistent changes to uncertainties or Wiener weights.

## Results

### The need to deposit in reciprocal space

We recommend depositing results of cryo-EM reconstruction in reciprocal space before Wiener filter is applied to create the final map. Such deposit would consist of complex structure factors and their uncertainties for at least three maps, final reconstruction map and its two half maps, and Wiener filter coefficients that could be common for all three maps.

This creates several advantages:

1. in cryo-EM SPR, information is accumulated in reciprocal space, so depositing in reciprocal space enables for proper SNR estimation that depends on the number of contributors in a particular position of reciprocal space.
2. the parameters calculated during reconstruction: the measured signal, its sigma *σ*, and the Wiener weight for each reciprocal space point can be used in downstream programs that refine in reciprocal space^11, 12^.

One would expect substantial improvement in atomic model refinement when using reciprocal space data. Optimal refinement depends on accurate sigma *σ* estimates, and the refinement target function should not use Wiener-filtered data. However, deposited real-space maps do not include variable *σ*, and the final maps are Wiener-weighted. Wiener filtering introduces strong resolution-dependent modulation of the structure factors, so when it is used with a target function that estimates B-factors, which are themselves resolution-dependent, the resulting B-factor values become severely overestimated. Servalcat preforms atomic model refinement in reciprocal space^11, 12^, so it could immediately take advantage of the type of deposition we propose. Other programs, such as Phenix^3, 13^, could adjust their refinement approaches to handle this additional information. Currently when only real space maps can be used as input, Servalcat works well for cases with isotropic coverage using non-Wiener-filtered half maps.

Depositing separate files for non-Wiener filtered data and the corresponding Wiener filter coefficients would also facilitate the calculation of difference maps. In generating a difference map between a reconstruction and its atomic model, the structure factors of the model are subtracted from the unweighted (non-Wiener-filtered) structure factors of the reconstruction. After computing these differences, the Wiener filter coefficients are applied to obtain a filtered difference map suitable for visualization. The resulting map closely approximates the log-likelihood gradient map.

Data deposited in reciprocal space can also make data storage on the EMDB server more efficient, as there is no need to deposit data past the effective resolution limit. Currently real space maps are often oversampled, with pixel sizes smaller than half of the resolution limit, because oversampled maps are more visually interpretable. If deposited in reciprocal space, real space maps can still be calculated with pixel size about 1/3 or 1/5 of the resolution limit on the user side, similar to what is done in crystallography. Sharpening can also be applied on the fly during map calculation using such reciprocal space data.

One important distinction from crystallographic practice concerns the calculation of R-free-like statistics. In cryo-EM, it is inappropriate to compute traditional R factors because the quantities refined are complex structure factors that include both amplitude and phase information, whereas R factors are defined solely on amplitudes. Although R-free serves as a hold-out statistical test in crystallography, defining an equivalent measure in SPR is not straightforward^14^. In cryo-EM SPR, points in reciprocal space are correlated because the 3D particle image is analyzed within a box volume much larger than the particle’s extent (Figure 4).

However, due to the absence of the phase problem, we have twice as much data available for refinement, which makes refinement bias weaker. An alternative to a hold-out test is to calculate unbiased refinement statistics based on random data perturbation followed by re-refinement. Comparing the structure factors of perturbed data and those calculated from a model refined against perturbed data can estimate resolution-dependent bias coefficients in this refinement. These coefficients can then be applied to scale up the disagreements between the model and the data, generating unbiased statistics equivalent to a knife-edge test such as R-free. Another variant of this approach is to perform two independent atomic model refinements using half datasets and then compare the model structure factors with the other half datasets not used in that model refinement (Figure 5). Debiasing through perturbation analysis has been implemented in the program DM, though in a somewhat different context^15, 16^.

**Figure 5.**
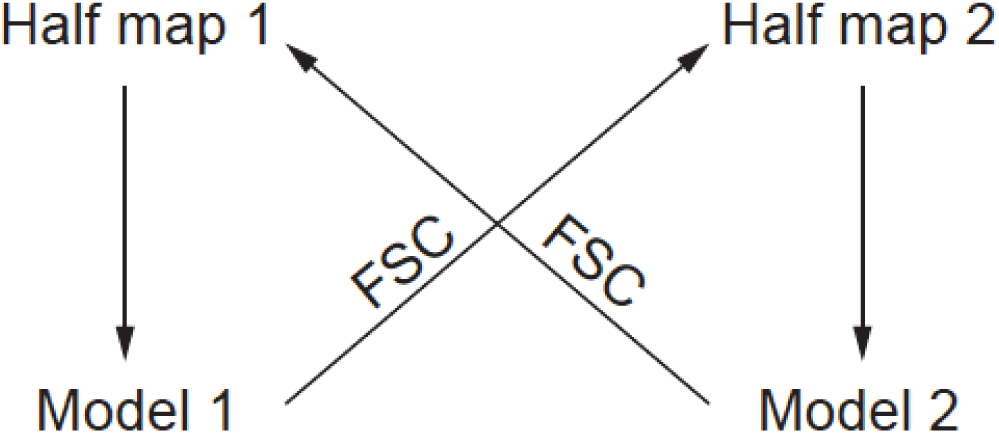
Two independent atomic model refinements using half datasets can be performed to obtain unbiased agreement statistics between the model and the map. Model 1 is refined against half map 1, and Model 2 is refined against half map 2. Independent FSCs can be calculated between Model 1 and half map 2 and Model 2 and half map 1. The expected FSC can then be calculated by converting two FSCs to SNRs, adding them in squares and converting the sum back to expected unbiased FSC. The final model should be refined against the final map weighted by statistics based on unbiased FSC.

### Masks as a real space filter to modify variance information from reciprocal space

We also recommend making the deposition of the molecular mask used in half-maps FSC calculations mandatory and advocate discontinuing the use of “unmasked” FSC for PDB validation, as well as reducing its use in general. First, calculating the half-map FSC without an appropriate explicit or implicit mask does not accurately reflect the true resolution of the reconstruction. Second, the “unmasked” calculation is not uniquely defined, any modification applied in real space can effectively act as a mask. This is particularly relevant for anti-aliasing filters and related reconstruction kernels. Depositing the actual mask, or the mask volume in the case of implicit masking, would enable the PDB to validate masked FSC results reported by the depositor. An example of a non-obvious mask occurs in membrane protein reconstructions, where masking out micelles surrounding the transmembrane region can significantly improve the apparent resolution by reducing refinement bias and removing noise contributed by the micelle volume. The issue of depositing mask-related information for methods such as cryoSPARC’s non-uniform refinement, or similar algorithms that employ combined reciprocal-space and real-space filtering, should be addressed separately.

One of the reasons for calculating unmasked half-map FSC arose from the fear of bias that can be introduced by masking^17^. This concern is related to the question of how a mask as a top-hat function correlates with the actual structure of the macromolecule. This problem has been addressed in crystallography through the analysis of the Babinet’s effect^18–20^. At lower resolution (∼10 Å), the correlation between the mask and the protein’s scattering is strong, so masking could introduce bias. At higher resolution (∼5 Å or better), the correlation rapidly approaches zero^20^, so the mask primarily functions as a noise filter and does not introduce significant bias after convergence (Figure 6). An incorrect mask may slow down convergence, but that is a separate problem not related to the final FSC being overestimated.

**Figure 6.**
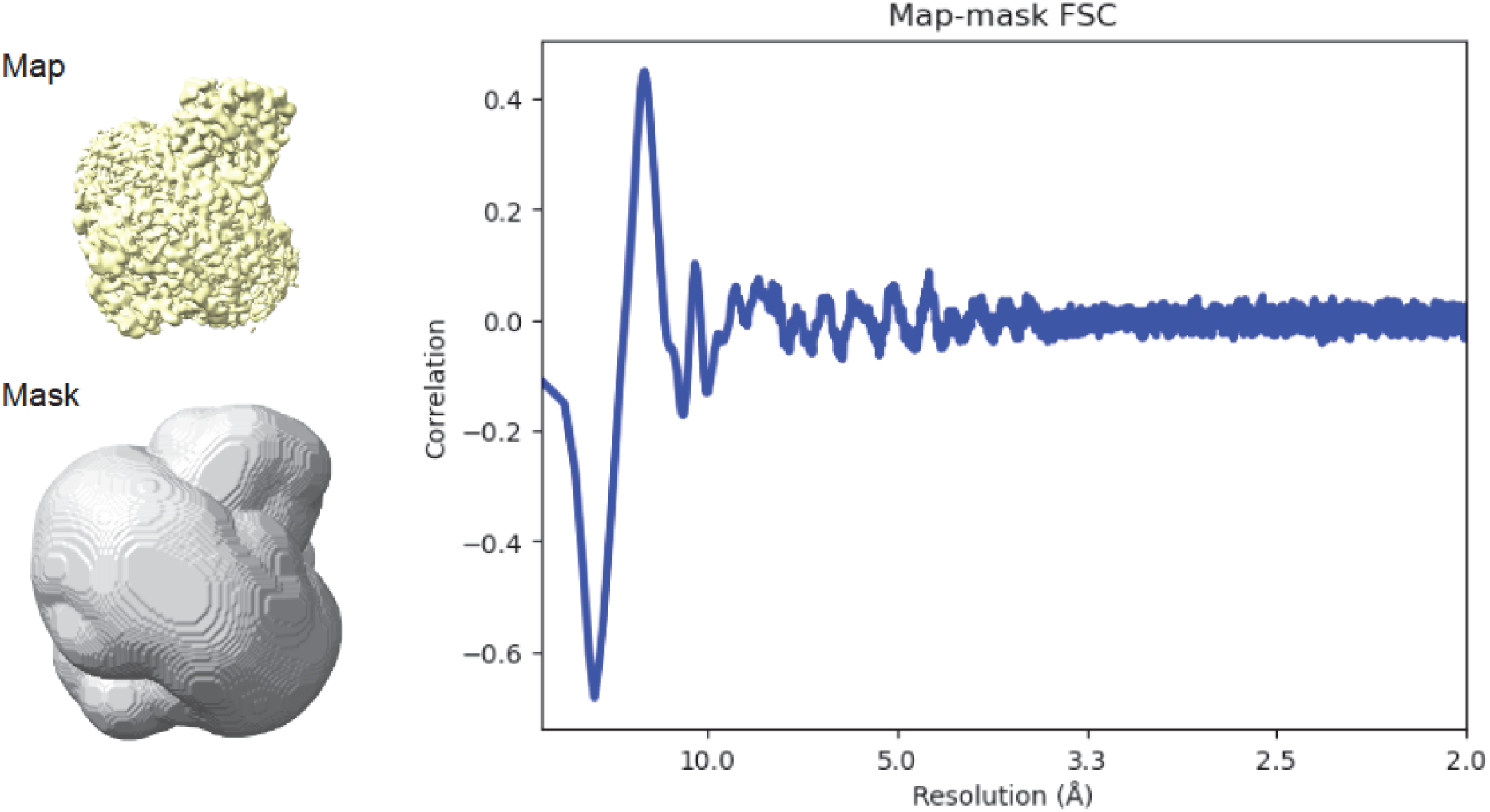
The molecular mask is not correlated with the model at resolution higher than 4 Å and thus the use of such masks cannot bias the FSC at high resolution. The EMD-28877^24^ is used as an example with the map-mask FSC calculated with Phenix^3^. Concerns about masking-induced bias result from historical experiences with lower resolution reconstructions, where the mask and model were strongly (anti)correlated at ∼10 Å. See also Figure 5 in Kostrewa^20^, where this dependence is represented as values of phase differences approaching 180° due to solvent and protein occupying mutually exclusive volumes.

Masks that do not cut into protein density cannot artificially increase the apparent resolution of the data, as they do not have an effect in modifying the signal from the protein.

If no molecular mask is provided, the calculation of “unmasked” FSC should employ a spherical mask with a diameter equal to the reconstruction box length, rather than a cubic mask that spans the full reconstruction box size. This is because particles could be diagonally aligned to the box, causing the corners of the particle box to project outside the reconstruction box during back projection (Figure 4). Due to the periodic boundary condition inherent in FFT, any signal extending outside the box will wrap around and reappear elsewhere inside the box, an aliasing artifact. Failing to apply a circular mask on the particle can also introduce other artifacts, such as noise accumulation in the corners of the reconstruction box. One should also note that some reconstruction algorithms, such as those using Kaiser-Bessel interpolation^21^, inherently apply an implicit, spherically symmetric mask during reconstruction.

### Data with preferred orientation

In cases of strong preferred orientation, where large portions of the reciprocal space are undersampled, calculating the FSC or defining a resolution limit based on it becomes meaningless, as it averages signals of uneven directional quality. This situation is similar, but not identical, to anisotropic diffraction and more closely resembles the missing cone problem. At present, there is no statistical descriptor of information content for such cases, indicating a clear gap in current methodologies. In both crystallography and cryo-EM, resolution is defined by the volume of reciprocal space containing useful signal that contributes to the map. The situation becomes more complex when this volume is not spherical. To address this, reciprocal-space volumes of arbitrary shape can be approximated by spheres of equivalent volume, using the resolution of that sphere as a single-number descriptor. This value represents the total amount of information in reciprocal space and provides the simplest conceptual descriptor for directionally non-uniform resolution. To achieve this, we propose calculating an effective resolution by summing the Wiener weights across reciprocal space and then reporting the resolution cutoff at which the number of reciprocal space points equals the sum of the Wiener weights. The same calculation can also be used in crystallography.

### A complete list of deposition requirements necessary to preserve information from reconstruction

To ensure that deposited cryo-EM data fully preserve the experimental information and can be independently validated or reanalyzed, we propose that depositions include a complete reciprocal-space description of the reconstruction. Each deposition should contain the original reconstruction in reciprocal space for the entire molecular assembly, or, when local refinement or motion decomposition has been performed, a set of reconstructions corresponding to those regions or particle subsets. The associated half reconstructions should also be included to enable independent validation through half-map FSC calculations that reflect the true weighting and directional sampling of the data. In addition, the molecular masks used during refinement and FSC calculation must be deposited. These masks, whether explicit or implicit, provide essential context for evaluating resolution and ensuring that results can be reproduced and compared across software packages.

The Crystallographic Information File (CIF) format^22^, a restricted derivative of the Self-Defining Text Archive and Retrieval (STAR) format^22^ used in cryo-EM, already provides the flexibility needed for such depositions^23^. All reciprocal-space reconstructions can be included in a single CIF file, with multiple values stored for each Miller index. This approach maintains compatibility with established databases and analysis tools, while supporting cryo-EM-specific information such as local weighting factors, anisotropy parameters, and per-point uncertainty estimates. Such depositions can also be made more compact by avoiding both spatial and reciprocal space oversampling. Optional real-space data can also be deposited for convenience and presentation. For instance, composite or enhanced real-space maps, such as AI-processed or filtered versions, may assist in visualization and model building (See Supplementary Information), but they should be recognized as derivative products that do not add information relevant to atomic refinement or validation. This deposition framework preserves critical statistical and structural details from modern cryo-EM reconstructions while remaining compatible with current infrastructure. By shifting the focus from real-space maps to reciprocal-space data, it ensures that deposited datasets remain complete, interpretable, and suitable for independent validation and future methodological advances.

## Discussion and Conclusions

Deposition of data to the PDB and EMDB is the important final step in the cryo-EM structure determination process and should preserve all information necessary to accurately assess the quality of the structural solution and enable downstream analyses. Although the current guidelines aim to reduce bias, improve comparability between datasets, and prevent unsavory manipulations, they do not fully account for the capabilities and practices of modern cryo-EM data processing software. Consequently, these standards can introduce ambiguity in data quality assessment, lead to information loss, particularly in challenging cases such as structures affected by strong preferred orientation, and create confusion about what should be deposited and how model quality should be interpreted (see Supplementary Information).

To address these limitations, we propose an alternative, complementary approach to the current standard. We recommend: (1) depositing cryo-EM reconstruction results in reciprocal space, (2) including explicit or implicit molecular masks as a mandatory part of the deposition, and (3) removing restrictions on filtering. Depositing data in reciprocal space preserves direction-dependent, non-Wiener-weighted signal and associated uncertainties for each reciprocal-space point. This provides a more accurate representation of orientation coverage and significantly benefits downstream atomic model refinement. Furthermore, half-maps FSC can still be calculated and validated using this information. Including molecular masks in the deposition improves the interpretability of FSC curves, generates more meaningful resolution estimates, and supports reproducibility.

Resolution normalization of data and uncertainties remains a complex and not fully addressed subject in cryo-EM. Because of the complex behavior of lenses and detectors, the signal contributing to the final reconstruction has already been through multiple levels of resolution-dependent manipulations. Directional motions further add variability, both directional and resolution dependent, to the maps. While directional effects are generally averaged out during reconstruction, the resolution-dependent components persist. All of these add to the complexity in how signal and uncertainty can be modeled.

The deposition standards proposed here, which include sigma values for each reciprocal-space point, do not fully resolve all aspects of modelling uncertainty. However, they preserve information that can be effectively described by existing software and provide a foundation for future tools to reanalyze refinements and improve uncertainty modeling.

## Author Contributions

Author contributions were as follows: Z. Otwinowski, Y. Guo, R. Bromberg and D. Borek analyzed the problem and wrote the manuscript.

## Supplementary Materials

### Case 1: Discussion about real space filtering

https://www.jiscmail.ac.uk/cgi-bin/wa-jisc.exe?A2=ind2508CL=CCPEMCP=34767 “I have rarely (never?) seen an unfiltered map deposited as the main map. The complication that Guillaume might have referred to here is that there are now machine-learning-driven software tools that produce improved maps (DeepEMhancer being one of them, but there are more and newer ones too - EMReady seems popular). The last time this issue was discussed, the majority expert opinion was that such “emhanced” maps should not be used for coordinate refinement. But they might be used to make figures, leading to the situation that the refinement map and figure map might not be the same map. This is not entirely new - we sometimes use different filtering resolution/sharpening levels to visualise poorly ordered peripheral regions in figures, which leads to a similar situation.

I believe that Guillaume’s suggestion is to deposit the conventionally post-processed map as the main map (as this is usually/hopefully used to refine the coordinates) and the de-noised/enhanced/otherwise “manipulated” map as an additional map. At the risk of stating the obvious: I agree with this suggestion.”

### Case 2: Discussion about model to map cross correlation

https://www.jiscmail.ac.uk/cgi-bin/wa-jisc.exe?A2=ind2009CL=CCPEMCP=R12021

“The model vs. data CC is only 0.73, whereas the CC is 0.84 when I refine against the Relion-postprocessed map. CC is a function of sharpening in real space – any type of noise filtering outside the mask will improve CC, we should use masked data to see whether enhanced map (anything past Wiener filter and done in real space, including later AI maps) produces better refinement results. – Model to Map CC is a good measurement for unenhanced map quality, but not a good measure for model quality. Improvement in CC does not always indicate improvement in model.

Map enhancement does not provide additional information over the original reciprocal space information, but makes the map in real space more interpretable. Not just the EMhancer maps, but any real space maps provided by cryoEM software.

Reciprocal space has structural factor and sigma --- reciprocal space to real space map is a set of procedures involve variable weighting (sharpening). One can create a blend of the sharpening for different regions in real space to create an enhanced real space map. Such sharpening has no impact on refinement in reciprocal space because it is absorbed in weights (but will have impact in real space refinement).”

These two discussions show community concerns relevant to the issues presented in the main text.

Due to Fourier localization theorem, flattening a molecular map outside the mask does not affect the refined result, <u>except for statistics based on correlation coefficient (CC) between</u> <u>the model and the map</u>. Therefore, improvements in CC must be interpreted with caution. An increase in CC for a filtered map can occur without any real improvement in the model, for example, by filtering noise outside the molecular model. Additionally, more complex real space procedures modifying maps resulting from cryo-EM SPR may have features of the model building. For instance, alpha helical density can be modified to follow more closely an ideal alpha helix. This can again increase the CC between the modified map and the model, without any improvement in the refinement target. However, it has large value in visualization and initial model building. For this reason, any additional real-space-modified maps used during model building or inspection should also be deposited, and their deposition should be mandatory. These post-processed maps should be used only for visualization, not as targets in refinement.

